# Mapping biology from mouse to man using transfer learning

**DOI:** 10.1101/2019.12.26.888842

**Authors:** Patrick S. Stumpf, Doris Du, Haruka Imanishi, Yuya Kunisaki, Yuichiro Semba, Timothy Noble, Rosanna C.G. Smith, Matthew Rose-Zerili, Jonathan J. West, Richard O.C. Oreffo, Mahesan Niranjan, Koichi Akashi, Fumio Arai, Ben D. MacArthur

## Abstract

Biomedical research often involves conducting experiments on model organisms in the anticipation that the biology learnt from these experiments will transfer to the human. Yet, it is commonly the case that biology does not transfer effectively, often for unknown reasons. Despite its importance to translational research this transfer process is not currently rigorously quantified. Here, we show that transfer learning – the branch of machine learning that concerns passing information from one domain to another – can be used to efficiently map biology from mouse to man, using the bone marrow (BM) as a representative example of a complex tissue. We first trained an artificial neural network (ANN) to accurately recognize various different cell types in mouse BM using data obtained from single-cell RNA-sequencing (scRNA-Seq) experiments. We found that this ANN, trained exclusively on mouse data, was able to identify individual human cells obtained from comparable scRNA-Seq experiments of human BM with 83% overall accuracy. However, while some human cell types were easily identified, others were not, indicating important differences in biology. To obtain a more accurate map of the human BM we then retrained the mouse ANN using scRNA-Seq data from a limited sample of human BM cells. Typically, less than 10 human cells of a given type were needed to accurately learn its representation in the updated model. In some cases, human cell identities could be inferred directly from the mouse ANN without retraining, via a process of biologically-guided zero-shot learning. These results show how machine learning can be used to reconstruct complex biology from limited data and have broad implications for biomedical research.

---

The translational biomedical research pipeline typically consists of a sequence of phases that starts with a discovery phase, which usually involves experiments on cell lines cultured *in vitro* as well as *in vivo* studies in model organisms, and ends with carefully controlled clinical, review and monitoring phases.^1^ The eventual success of this pipeline depends upon effective transfer of information from one phase of the process to the next. Despite the tremendous cost associated with translational research failure,^2^ this information transfer process remains poorly understood.

Transfer learning (TL) is the branch of machine learning that takes information derived from one setting and applies it to improve generalization in another area.^3^ The basic idea of TL is to mimic the human ability to learn new concepts from limited examples by associating new information with prior understanding. In the TL process, information gained from solving a problem in a “source” domain is passed to another related problem in a “target” domain thereby improving target domain performance. The gain from such knowledge-transfer is particularly apparent whenever data is abundant in the source domain but scarce in the target domain. In this case, new concepts can be effectively learnt in the target domain from very few training samples, via leveraging of prior knowledge.

Here, we show how TL can be used to map bone marrow (BM) biology from mouse to man. The problem of passing information from a model organism (the source domain, here the mouse) to the human (the target domain) was chosen because it is central to successful translational research. Bone marrow was chosen because it is a complex tissue, consisting of numerous different cell types, present in differing proportions, with a well-established physiology in mouse that is broadly conserved, and yet only partially understood, in human.

## Mapping mouse bone marrow

To begin, we collected gene expression signatures using droplet-based scRNA-Seq (Drop-Seq^4^) from unfractionated total bone marrow (TBM) samples as well as from weakly lineage-depleted bone marrow (DBM) Cd45/Ter119 dual negative subsets in order to enrich for rarer cell types, from three different mice (Fig. 1a). Overall, 6,800 single-cell transcriptomes were sequenced, yielding greater than 9×10^4^ reads per cell on average. Following pre-processing and filtering, a total of 5,504 cells were retained, expressing on average 2,684 transcripts per cell.

**Figure 1.**
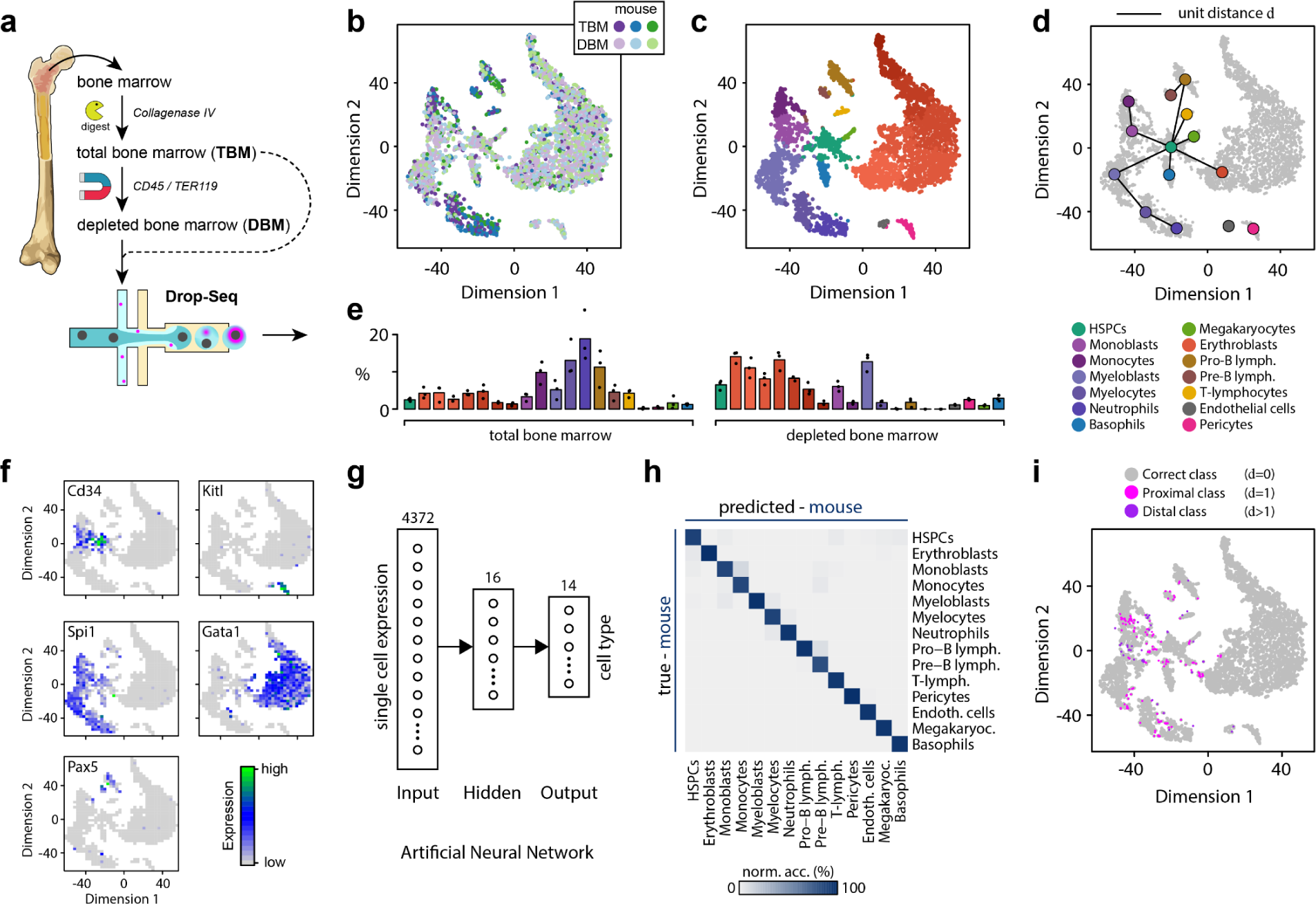
Dissecting the cellular heterogeneity of the mouse BM. **(a)** Experiment schematic. Single-cell RNA-sequencing was performed on total and depleted (CD45-/Ter119-) BM cells. **(b)** Projection of data onto two dimensions using t-distributed stochastic neighbor embedding (tSNE^8^) highlights the BM population structure and overlap between biological replicates (*n*=3). **(c)** Cell types were identified using unsupervised clustering followed by annotation of clusters according to localization of known markers for different cell types. **(d)** Clusters naturally arrange in accordance with the known BM lineage tree. The lineage tree shown is taken from Ref. ^9^. HSPCs: hematopoietic stem and progenitor cells **(e)** Relative abundance of cell types in total and depleted bone marrow samples. Bar height indicates the mean over the biological replicates (*n*=3). **(f)** Key markers of the main branches of the hematopoietic lineage tree and niche cells localize to distinct clusters in the data. The following representative markers are shown: stem and progenitor cells: Cd34; niche cells: Kitl; myeloid lineage: Spi1; erythroid lineage: Gata1; lymphoid lineage: Pax5. See Fig. S1 for localization patterns of a range of other markers. **(g)** Schematic of the artificial neural network (ANN) used to identify cell types from gene expression profiles obtained from mouse BM cell samples. The ANN consists of an input layer, a 16-node hidden layer, and a 14-class softmax output layer. **(h)** Confusion matrix of validation data, showing accurate classification of cell identities by the ANN. Data displayed is the average over a 5-fold cross-validation. **(i)** Distribution of misclassified cells in the training data. Color represents the distance *d* between the true and predicted label in the cell lineage tree in panel e.

We then performed unsupervised clustering to identify the various hematopoietic and niche-cell types present (see **Methods**). Despite the significant technical variability that is typically encountered in scRNA-seq data^5^ we found that cells clustered according to their type, rather than the mouse from which they were obtained, suggesting the presence of a common and robust “map” of the mouse bone marrow (Fig. 1b-d and Fig. S1, Fig. S3a). Assignment of cell identities to clusters was performed by examining the localization of established lineage markers to distinct clusters (see Fig. 1f, Fig. S1 and **Methods**). Our cluster annotation was in accordance with other recent publications.^6,7^ In total, we identified 19 cell populations, covering the erythroid, myeloid and lymphoid branches of hematopoietic lineage tree, as well as separate populations of non-hematopoietic supporting cell types including endothelial cells and pericytes (Fig. 1c-f).

Three features of this clustering are notable. Firstly, the proportion of cells in each cluster varied considerably, reflecting the balance of different cell types present in the mouse BM (Fig. 1e). Clusters associated with rare cell types, such as hematopoietic stem and progenitor cells (HSPCs), contained very few cells. In contrast, clusters associated with abundant cell types, such as erythroid cells, contained large numbers of cells. To gain resolution on rare/immature cell types the depletion protocol we used reduced the relative abundance of various mature cell types in the DBM fraction – including monocytes (−8.1 ±3.2% relative to TBM), myelocytes (−11.4 ±5.6% relative to TBM), pro-B-lymphocytes (−9.4 ±3.8% relative to TBM), while neutrophils, pre-B- and T-lymphocytes were completely ablated – while enriching for immature cells including HSPCs (+4.0 ±0.9% relative to TBM), myeloblasts (+7.5 ±3.7% relative to TBM), monoblasts (+2.8 ±1.7% relative to TBM), erythroblasts (+5.4 ±1.9% relative to TBM), Pericytes (+0.9 ±0.3% relative to TBM) and endothelial cells (+2.1 ±0.5% relative to TBM).

Secondly, while some clusters represent distinct cell identities, others are associated with the discretization of continuous maturation processes. For example, the erythroblast cluster (shown in red in Fig. 1c) consists of a heterogeneous mixture of cells at different stages in the erythrocyte maturation process, representing a gradual transition from immature pro-normoblast to late normoblast (see Fig. S1). This observation will be important later, when we translate BM biology from mouse to man.

Thirdly, it is well-established that cell types in the hematopoietic cell lineages of the BM are arranged according to a hierarchical structure.^9–11^ By considering adjacency relationships between clusters, we were able to broadly recapitulate the known structure of this hierarchy, indicating that the clustering structure that we observed captures salient features of the mouse BM biology (Fig. 1d).

Collectively these clusters, and the spatial relationships between them, constitute a reference map of the mouse BM. However, this map is not in a form directly amenable to comparison with human BM. To do this, we trained an artificial neural network (ANN) to classify individual cells from their gene expression profiles. Since we ultimately wanted to compare this map with a similar map of human BM we restricted our analysis to those genes with a unique human orthologue (see **Methods**). Because cell identities were determined from unsupervised clustering of the data, this is an easy learning problem readily solved by a multi-layer perceptron with a single hidden layer of 14 units (Fig. 1g-i). The resulting model performed well, achieving average balanced classification accuracy of 96.7±0.9% (standard deviation over 5-fold cross-validation), and was able to reliably identify cells of every type (Fig. 1h).

Notably, misclassification was largely constrained to cells in proximity to cluster boundaries imposed along continuous developmental trajectories (Fig. 1i). To dissect this observation further we systematically investigated misclassification by taking advantage of the fact that the classes in the mouse training data are arranged according to a biologically meaningful hierarchy that encodes the BM lineage tree (see Fig. 1d). Whenever misclassification of a cell occurred, we determined the relationship between its true class and its (falsely) predicted class. We denoted a misclassification to be proximal if the predicted class is immediately adjacent to the true class in the lineage tree and distal otherwise. Overall, a low incidence of proximal misclassification was observed (2.7±0.5%; mean ± s.d., n=5), while distal misclassification occurred even more rarely (0.8±0.4%; mean ± s.d., n=5).

Moreover, patterns of misclassification were not uniform. For example, cells in the HSPC cluster were most likely to be misclassified (proximal: 9.9±2.3%; distal: 0.3±0.7%; mean±s.d., n=5). This is likely partly due to the limited number of HSPC training samples available. It also indicates that HSPCs represent a heterogeneous population with expression patterns that partially intersect with several other cell types. This observation agrees with recent studies in mouse, human and zebra fish, that have shown that the HSPC pool is a particularly variable cell population,^7,9,10,12,13^ and highlights the fact that classification accuracy will depend upon both the amount of data available for training and the heterogeneity intrinsic to the cell population being considered. This issue will also be important when we consider transferring mouse biology to the human.

To further dissect the biological basis of this classifier, we also conducted an information-theoretic sensitivity analysis designed to determine subsets of genes (i.e. predictors) that are most strongly associated with each cell identity (i.e. output classes, see **Methods** for details). This analysis recapitulated well-established molecular markers of BM cell type identities (see **Table S1**).

For instance, among the top-ranking features associated with the HSPC identity are *Angpt1* and *Myct1*, both known regulators of stem cell proliferation;^14,15^ *Irf8*, a monocyte lineage determinant,^16^ was most strongly associated with the monoblast identity; *Ccr2* and *Ctss*, which are known to play a central role in chemotaxis and antigen presentation of monocytes,^17,18^ were most influential for monocyte classification. Assignment of the myeloblast identity was highly sensitive to *Prtn3* (also known as Myeloblastin), while transcripts encoding components of secretory vesicles that are sequentially produced during myelopoiesis,^19^ and define the morphologically distinct stages of myeloblasts (primary granules; *Elane*), myelocytes (secondary granules; *Ltf*) and neutrophils (tertiary granules containing the neutrophil-collagenase *Mmp8*) also strongly influenced these class assignments. Similarly, various different immature lymphocytes were discriminated based on the expression of *Vpreb3* (pro-B-cells), *Cd74* (pre-B-cells) and *Ccl5* (TEM cells); while (peri-)vascular cells were determined based on *Serping1* (pericytes) and *Fabp4* (endothelial cells) expression among other genes. **Table S1** contains a complete list of all the top-ranking genes associated each cell type.

To investigate the functional significance of these gene sets we also performed Gene Ontology (GO) term analysis (see **Methods**). We found that significantly enriched GO terms associated with these gene lists summarized the biological function of their associated cell type. Key GO associations included *hemopoiesis* for HSPCs (*p*=8.5e-5; modified Fisher’s exact test), *blood coagulation* for megakaryocytes (*p*=1e-8), *B-cell receptor signaling* for pro-B- and pre-B-cells (p=1.2e-8; p=3.9e-10); *T-cell receptor signaling* for T-lymphocytes (p=1.6e-10); *cell adhesion* and *osteoblast differentiation* for pericytes (p=7.6e-10; p=9.5e-9); *cellular response to VEGF* for endothelial cells (p=8.4e-8); *positive regulation of mast cell degranulation* for basophils (p=1.2e-6); and *innate immune response* and related terms for monocyte- and granulocyte-lineages. **Table S2** contains a complete list of GO terms associated with each cell type.

Collectively, these results indicate that our ANN captures the essential biology of the mouse BM and can accurately discriminate between mouse BM cell types based upon differences in biologically significant gene expression patterns.

## Mapping human bone marrow

We next sought to determine the extent to which the biology learnt in the mouse “source” domain could be transferred to the human “target” domain of true interest. To do this, we sequenced BM samples from three patients, undergoing routine hip replacement surgery at Southampton General Hospital. In total, ~25,000 single-cell transcriptomes were sequenced yielding on average 5×10^4^ reads per cell. As with the mouse, we sequenced unfractionated BM as well as depleted populations in order to enrich for rarer cell types. Following pre-processing and filtering of low-quality cells (see **Methods**) we obtained data for 9,394 cells expressing on average 3,070 transcripts per cell. As with the mouse data we then performed unsupervised clustering to identify the various hematopoietic and niche-cell types present and assigned cell identities based upon localization of established lineage markers (see Fig. S2 and **Supplementary Material**). As with the mouse data this analysis resulted in a set of single cell transcriptomes in which each cell is annotated with a unique identity determined by unsupervised clustering.

We subsequently assessed the extent to which our mouse classifier, which was trained exclusively on mouse data, was able to predict human cell identities (Fig. 2a). We found that the mouse classifier predicted human cell identities remarkably well, achieving an average balanced accuracy of 83.3%. Notably, this overall accuracy was not consistent across all cell classes: rather, accuracy ranged from 60.0 to 98.0% for individual cell classes (Fig. 2b). Thus, while some human cell types were identified remarkably well by the mouse classifier, indicating strongly shared biology, others cell types were much more poorly aligned, indicating systematic differences in underlying biology between the species. For example, human erythroblasts and T-lymphocytes were rarely misclassified by the naïve mouse model (which achieved 97.5% and 98.0% balanced accuracy in identifying these classes respectively), while other cell types were frequently misclassified (Fig. 2b).

**Figure 2.**
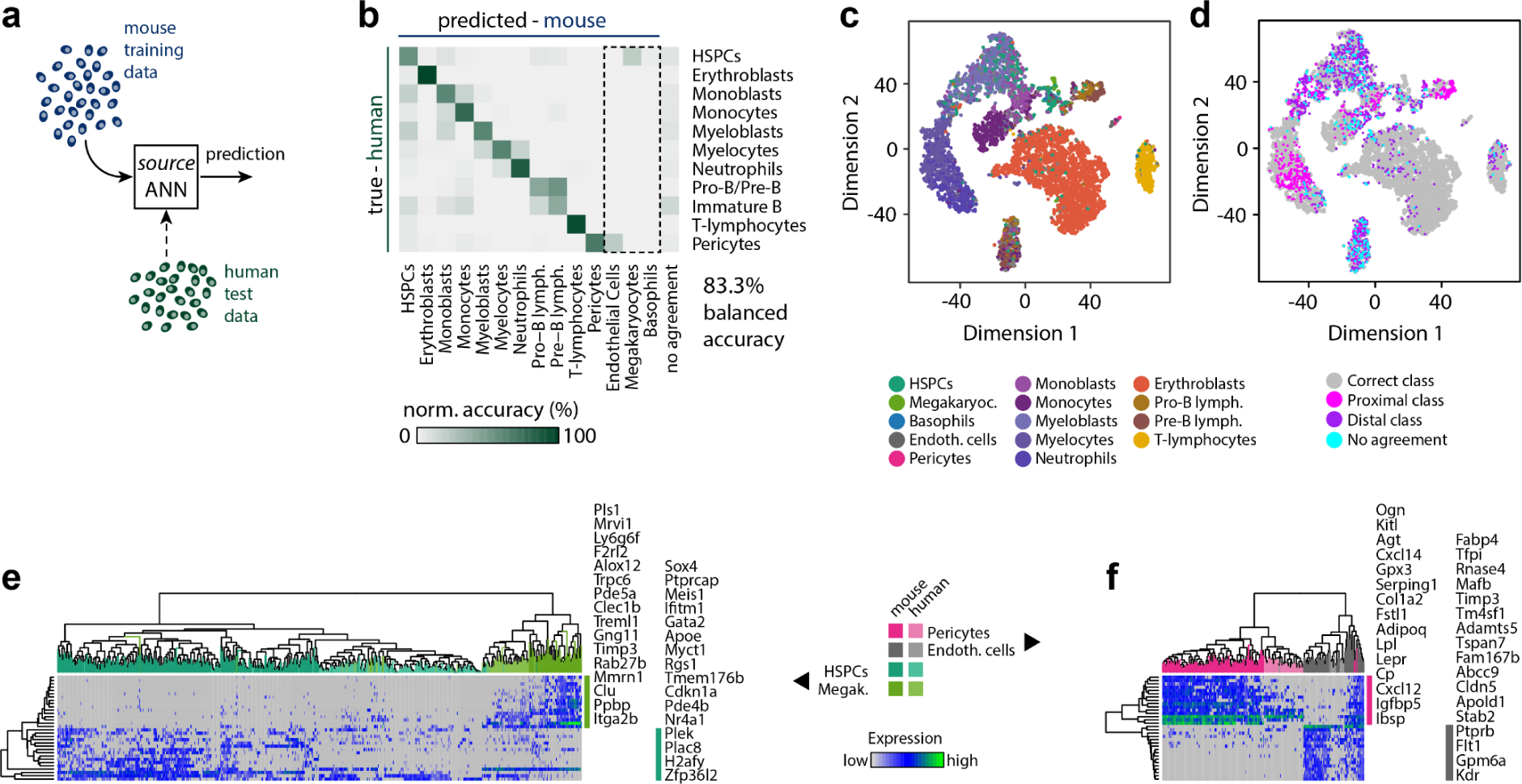
Bone marrow biology maps partially from mouse to man. **(a)** Schematic of the naïve transfer process. ANN trained in the source domain (mouse) is used to classify test data from the target domain (human). (**b)** Confusion matrix of classification consensus from 5-fold cross validation. The dashed box highlights cell types identified in the mouse but not the human data. **(c-d)** Projection of human data onto two dimensions using tSNE^8^. Points represent cells coloured by (c) predicted cell identity or (d) misclassification. **(e-f)** Co-clustering of expression patterns in mouse and human cells discriminates human HSPCs from megakaryocytes (e) and pericytes from endothelial cells (f). In both panels clustering is performed using the top-ranking genes from sensitivity analysis

As with the mouse data, we found that misclassification of human cells was commonly proximal in nature (13.9% versus 8% for distal misclassification overall), suggesting that the mouse classifier had partially learnt human cellular identities, and misclassification was not entirely artefactual (Fig. 2b-d). For instance, human HSPCs were systematically misclassified as one of their proximal descendent classes (17.8% misclassification), but less frequently distally misclassified (6.9% misclassification). A similar pattern of misclassification of mouse HSPCs was also seen (Fig. 1i). Likewise, myelocytes were systematically misclassified as their progenitors, myeloblasts, or their descendants, neutrophils (29.6% proximal versus 8.7% distal misclassification) (Fig. 2b-d). However, this pattern was not universal.

For example, while human immature-B lymphocytes were commonly misclassified as their progenitors, pro-B and pre-B lymphocytes, they were also systematically misclassified as a range of other types of progenitors (14.1% proximal misclassification versus 33.8% distal; see Figs. 2b-d), indicating that the mouse classifier was not able to fully resolve the human immature-B cell identity. Notably, the mouse data did not contain a comparable immature-B cell cluster, and so the mouse model was never explicitly trained to recognize the expression signatures of B-lymphocytes. Nevertheless, the mouse classifier assigned the majority of immature B lymphocytes to adjacent clusters and hence to the correct branch of B-cell development.

Taken together, these results indicate that while much biology is conserved between the mouse and human BM, there are systematic differences. These differences are important because they indicate the circumstances in which the mouse is likely to be a good model of human biology and when it will likely not; and they highlight instances where a comparison is not immediately possible.

## Discovery of hidden cell identities using zero-shot learning

While the mouse and human datasets contain data from many of the same cell types, some cell types were not resolvable in the human samples with accuracy comparable to the mouse. In particular, we could not identify distinguishable clusters associated with endothelial cells or megakaryocytes, yet both of these cell populations were clearly apparent in the mouse data. Because many aspects of BM are conserved between mouse and man, we next sought to determine if the mouse model could be used to help resolve the biology such hard-to-identify human cell types.

Interestingly, a significant subset (19.4%) of human HSPCs were identified as megakaryocytes by the mouse classifier (Fig. 2b, c). This high overlap is notable because HSPCs and megakaryocytes are proximal in the mouse hematopoietic cell lineage map (Fig. 1d), reflecting the fact that megakaryocytes emerge directly via differentiation from HSPCs^9,20,21^. This result suggested to us that the mouse classifier might be revealing aspects of human HSPC/megakaryocyte biology that are not apparent from unsupervised clustering of the human dataset alone. To investigate these differences further, we conducted sensitivity analysis (see **Methods**) to identify the genes that carry the most discriminatory information in distinguishing between megakaryocytes and HSPCs in the mouse classifier. Examination of co-expression patterns of these genes in human and mouse cells confirmed that megakaryocytes are characterized by broadly similar expression signatures in both mouse and human, and are distinguishable from HPSCs based on these expression patterns (Fig. 2f).

Notably, both mouse and human megakaryocytes expressed high levels of genes involved in platelet biogenesis such as *Rab27*b,^22^ *Ppbp*,^23^ and in platelet function (hemostasis) such as *Itga2*^24^ (encoding collagen receptor CD49b) and *F2rl2*^25^ (encoding coagulation factor 2) (Fig. 2f). Similarly, HSPCs in both species shared expression of key transcription factors such as Zfp36l2 and Sox4 (Fig. 2f) that are known to control stem cell self-renewal^15,26^.

Similarly, when shown to the mouse classifier a significant subset (20.3%) of human pericytes were identified as endothelial cells (Fig. 2b, c). While the ontogeny of pericytes and endothelial cells in the adult bone marrow remains unclear,^27^ both cell types are constituents of the vasculature, and are in close spatial proximity in the BM, again suggesting that the mouse classifier might be revealing aspects of human biology that are not apparent from unsupervised analysis of the human dataset alone. To investigate these differences further we again conducted sensitivity analysis (see **Methods**) to identify the genes that carry the most discriminatory information in distinguishing between endothelial cells and pericytes in the mouse model. Among the genes that were identified were a number of important endocrine modulators and sensors of energy homeostasis such as *Igfbp5* and *Lepr;*^28,29^ paracrine signaling molecules such as *Cxcl12*;^30^ and components of the iron cycle such as *Cp.* Examination of co-expression patterns of these genes revealed a substantial overlap between mouse and human pericyte expression patterns, indicating that much of the central molecular machinery of these cells is evolutionarily conserved (Fig. 2e). Similarly, both human and mouse endothelial cells shared expression of known angiogenic-signal receptors such as *Kdr*^31^ and the Vegf target gene *Fabp4*^32^ (Fig. 2e), again highlighting shared biology. However, we also observed that a subset of human pericytes clustered with mouse endothelial cells again suggesting that the mouse classifier was able to reveal aspects of human cell identities not apparent from the human data alone (Fig. 2e).

Collectively these results indicate that once encoded in a machine learning model, mouse data can be used to contextualize human data, identify evolutionarily conserved gene expression patterns and thereby provide insight into poorly resolved cell populations. In the machine learning literature, the process of object identification without training examples is known as zero-shot learning, and typically relies on importing prior knowledge from a related source domain.^33^ Here, because the mouse classifier encodes evolutionary conserved information, it can be used, in conjunction with prior knowledge of the BM lineage tree, to infer poorly resolved human cell populations via a process of biologically-guided zero-shot learning.

## Transferring biology from mouse to man

Since the mouse classifier was not able to accurately identify all human cell identities, yet appeared to be capturing aspects of evolutionarily conserved biology we next sought to determine if it could be used to train a more accurate model of the human BM. To achieve this objective, we re-trained the mouse classifier using a limited set of human BM cell gene expression signatures as additional training data (Fig. 3a).

**Figure 3.**
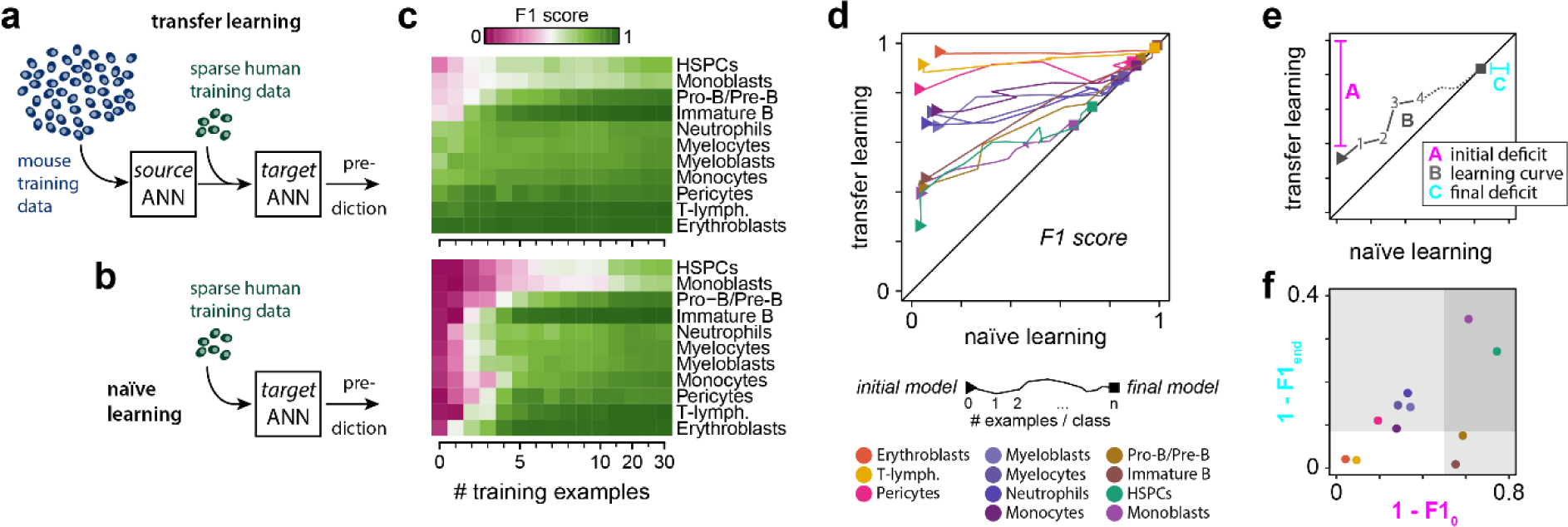
Mapping biology from mouse to man using transfer learning. **(a)** Schematic of the transfer learning process. Abundant data from source domain (here the mouse) is used to train a source ANN. Sparse data from the target domain (here the human) is used to fine-tune the parameters of the source ANN, thereby transferring knowledge from source to target domain. **(b)** Schematic of naïve learning as a control for transfer learning. Rather than updating the pre-trained mouse model, a series of separate ANNs are trained from random initial conditions on sparse data from the human target domain. **(c)** Both transfer and naïve learning improves with the number of human samples used for training (shown is data for 0,1, 2, …, 10, 15, 20, 25, 30 human cells per class). Transfer learning performance (top) and naïve learning (bottom). Displayed is the average F1 score from 5-fold cross validation as a measure of classifier performance. **(d)** Learning curves illustrate the evolution of classification performance starting from the initial mouse (triangle) to the final model (square; trained on 30 human examples per class). **(e)** Schematic to interpret the learning curves in panel d. Three features are of importance. A is the initial performance deficit = 1 – F1_0_, where F1_0_ is the F1 score of the mouse model in predicting human samples. B is the learning curve: each point on this curve plots the F1 score of the retrained mouse model against the naïve human model for a fixed number of human training examples from 0 to 30 per class. C is the final performance deficit = 1 – F1_end_, where F1_end_ is the F1 score of the naïve human model trained on 30 samples per class in predicting human samples. The line *y = x* is in black. On this line the naïve and retrained models have equivalent accuracy for the same number of human training samples. At this point the mouse biology has been forgotten, and equivalent learning can be achieved by a naïve human model. All learning trajectories eventually converge to this line. **(f)** Cell types may be grouped by their initial and final performance deficits.

We produced a series of ANNs by retraining the mouse classifier using increasing numbers of additional human training examples (Fig. 3a, c). Classification performance sharply increased on retraining, even when only a very small number of representative human training examples were used (Fig. 3c). Notably, classifier performance began to saturate when retraining using 4-8 additional human training examples for each class (44-88 cells in total). At this point retrained models achieved over 90% balanced accuracy (up from 83% in the naïve mouse model) and a significantly improved F1 score (88.4±0.8% up from 63.1±0.6%; mean ± s.d., n=5) indicating that human cell identities can be reliably predicted upon retraining with very few training examples (Fig. 3c).

In order to assess the extent to which pre-learning in the mouse source domain improves classification performance in the human target domain, an equivalent set of ANNs were trained without transfer from the mouse (i.e. from randomized initial conditions, referred to as naïve models; Fig. 3b-c). Since they benchmark the efficiency with which human BM biology can be learnt from low volumes of data without pre-training in the mouse these naïve models act as controls for the transfer learning process. To assess the information-transfer process we plotted classifier performance (here, the F1 score, which accounts for both the precision and recall of the classifiers) of transferred and naïve models against each other as the number of human training examples varied to produce characteristic learning curves (Fig. 3d, e). This analysis indicates the extent to which the biology of each cell type is shared between species (see Fig. 3d and further explanation in Fig. 3e). Four distinct groups of cell types can be distinguished based on their different characteristic learning curves (Fig. 3d).

The first group contains cell types with highly conserved phenotypes, which display high classification performance initially (i.e. the mouse classifier is able to identify human cells without additional training using human data) that does not improve significantly upon re-training in the target domain (i.e. using additional human data; Fig. 3d-f). This group includes erythroblasts and T-lymphocytes. These cell types: (1) are highly homogeneous in their expression patterns in human and thus are consistently classified; and (2) have a biology that is highly conserved between mouse and man. These cell types can be reliably identified from the mouse classifier and do not require any human training data to learn their representation. The mouse is a good model of human biology for these cell types.

The second group contains cell types that display good (but not excellent) classification performance initially, that does not improve significantly upon re-training with human data (Fig. 3d-f). The second group includes pericytes, myeloblasts, myelocytes, neutrophils and monocytes. These cell types: (1) are more heterogeneous in their expression patterns in human and thus are less consistently classified than group 1; and (2) have a biology that is highly conserved between mouse and man. This group of cell types requires a moderate amount of human training data to learn their representations.

The third group contains cell types that display initially low classification performance that improves rapidly upon re-training with human data (Fig. 3d-f). This group contains pro-B/pre-B lymphocytes, and immature B lymphocytes. These cell types: (1) are homogeneous in their expression patterns in human; yet (2) have a biology that is distinct between species. The biological differences between species for these cell types are in part due to differences in cluster definition in mouse and human data. Specifically, while the mouse ANN was trained to distinguish pro-B and pre-B cells, these cells are part of the same cluster in the human data. Hence, re-training involves separating the joint cluster of pro-B and pre-B lymphocytes from previously unseen immature B lymphocytes. This group of cell types requires a moderate amount of human training data to learn their representations.

Finally, the fourth group contains cell types that display low classification performance initially that does not improve significantly upon re-training with human data (Fig. 3d-f). This group contains monoblasts and HSPCs. These cell types: (1) are heterogeneous in their expression patterns in human; and (2) have a biology that is distinct between species. This group of cell types requires a large amount of human training data to learn their representations.

Collectively, this analysis shows how tools from transfer learning can be used to dissect those aspects of biology that will effectively transfer between the species and those aspects that do not.

## Discussion

Successful biomedical research is critically dependent on the effective transfer of information between different stages of the research pipeline. A critical step in this process is knowledge transfer from model organisms to the human. Here, we have shown how methods from transfer learning can be used to efficiently pass biological information between species using the bone marrow as an example. As increasingly detailed single cell maps of whole organism biology become more available, we anticipate that transfer learning approaches will provide essential tools for comparative physiology. More generally, the transfer learning philosophy is not limited to single cell data. Similar approaches are relevant whenever data is easily obtained from a data-rich source domain, yet hard to obtain in a related data-sparse target domain. As an example, understanding of rare disease biology may be substantially improved by leveraging understanding of related, yet common, diseases. Similarly, transfer learning could be used to pass knowledge of drug responses from pre-clinical model systems to the human. In conclusion we anticipate that transfer learning methods can be used to reconstruct complex biology from limited data and, will significantly streamline future biomedical research and help bring therapies more quickly and cost-effectively to the clinic.

## Methods

### Mouse tissue origin

Bone marrow from female 8-week old C57BL/6 mice was used in this study. All experimental work including mice were approved by the Kyushu University animal experiment committee.

### Human tissue origin

Excess marrow was collected from patients undergoing routine hip replacement surgery, with informed consent, and use of human tissue was approved by the regional ethics committee (reference 18/NW/0231).

### Bone marrow cell isolation

Mouse bone marrow mononuclear cells (mBM-MNCs) were prepared as described previously.^29^ Bone marrow was flushed from tibiae and femurs and digested with 1 mg/ml collagenase IV (Thermo Fisher, 17104019) and 2 mg/ml dispase (Gibco, 17105041) in Hank’s balanced salt solution (HBSS; Gibco, 14025092) for 30 min at 37 °C. Dissociated cells were treated with ammonium chloride solution to remove erythrocytes (155 mM NH_4_Cl, 12 mM NaHCO_3_ and 0.1 mM EDTA) for 5 min at room temperature, following 3x washes in HBSS.

Human bone marrow mononuclear cells (hBM-MNCs) were prepared described previously,^34^ with the additional removal of erythrocytes following density centrifugation through lysis in ammonium chloride solution (155 mM NH_4_Cl, 12 mM NaHCO_3_ and 0.1 mM EDTA) for 5 min at room temperature, following 3x washes in plain α-MEM.

### Magnetic cell sorting

Cells were immuno-labelled with magnetic microbeads for cell separation according to manufacturer’s instructions. Up to a total of 1×10^8^ BM-MNCs were used for each separation. Human cells expressing CD45 (Miltenyi Biotec, 130-045-801) or CD235a (Miltenyi Biotec, 130-050-501) and mouse cells expressing CD45 (Miltenyi Biotec, 130-052-301) or TER119 (Miltenyi Biotec, 130-049-901) were depleted using LS columns (Miltenyi Biotec, 130-042-401) according to manufacturer’s instructions.

### Collagenase release of bone lining cells from human bone marrow

Trabecular bone fragments obtained after the first step of cell isolation were incubated in 20 U/ml Collagenase IV (Thermo Fisher, 17104019) for 3h at 37°C under continuous rotation. Bone fragments were washed with plain α-MEM (Thermo Fisher, 12000-014) and cells released from ECM were filtered using a 40µm cell strainer.

### Single-cell RNA-sequencing

Single-cell sequencing was performed as described in detail elsewhere^4^ and alterations of the original protocol are reported below. Hydrophobic surface treatment of polydimethylsiloxane (PDMS) microfluidic devices was performed by incubating channels with 1% Trichloro(1H,1H,2H,2H-perfluoro-octyl)silane (Sigma-Aldrich, 448931) in Fluorinert FC-40 (Sigma-Aldrich, F9755) for 5-10min at RT. Syringe pumps to drive both aqueous and non-aqueous phases were made in-house according to published, open source protocols.^35^ Protocols for NGS library preparation described in Macosko et al. 2015 were closely followed and pre-amplification was conducted using 4+12 PCR cycles (95°C 3 minutes - 4 cycles of: 98°C 20s; 65°C 45s; 72°C 3min - 12 cycles of: 98°C 20s; 67°C 20s; 72°C 3min - 72°C 5min; 4°C hold). Processed libraries were sequenced using a NextSeq 500 system (Illumina) and NextSeq 500/550 High Output Kit v2 (Illumina, TG-160-2005).

### Sequence alignment

Sequence alignment was performed as detailed in Macosko et al. 2015 using the mm10 (GSE63472) and hg19 (GSM1629193) reference genomes and STAR (version 2.5.2b) for sequence alignment. Raw reads were demuliplexed and condensed into the digital gene expression matrix (DGE) using DropSeq tools (v1.0; Macosko et al. 2015), using a modified alignment score to reduce the number of reads discarded due to multiple alignment.

### Data pre-processing

Data was analyzed using the software R (version 3.5.0) and the Seurat package (version 2.3.1). A gene mapping between mouse and human was created using the orthologue annotation provided by Ensembl.^36^ Unscaled data was discretized (threshold >0) and the union of genes from both species previously identified as variable (threshold for mean 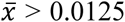, and 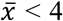; and log of dispersion > 0.5) in Seurat were selected for training if they were unambiguous orthologues.

### Cluster analysis

Clustering was performed in Seurat using the Louvain algorithm^37^ with resolution parameter set to 1.1.

### Cluster annotation

To assign meaningful labels to the clusters proposed by the Louvain algorithm, differentially expressed genes were identified using the likelihood-ratio test^38^ built-in to Seurat (with settings: prevalence > 25%; fold-change > 2; p-value < 0.001). Thus, obtained cluster markers were screened for previously described biomarkers for given bone marrow cell populations.

### Cell lineage tree

A qualitative description of the cell lineage tree was obtained from the literature.^9^

### Prerequisites to machine learning of cell identities across species

The problem of mapping between the cell identities of any number of species consists of two parts: Firstly, a mapping between features (here: genes), and secondly, a mapping between cell types. A useful feature map contains genes that were present in a last common ancestor. Such a mapping between features (orthologous genes in these data) can be easily obtained by adopting existing annotation from Ensembl that is based on sophisticated phylogenetic sequence analysis.^36^ In mouse and human only half of the genomic DNA can be mapped between genomes,^39^ limiting our analysis initially to the remaining conserved features. The other part, the non-conserved features, can thus be regarded separately from theses analyses and contribute foremost to the species-specific biology of cell types.

### Machine learning

Artificial neural networks (ANNs) were trained using the keras for R package (v2.2.4; https://keras.rstudio.com/) and the TensorFlow backend (v1.8.0; https://www.tensorflow.org/) on a GeForce GTX 1050 GPU (NVIDIA, Santa Clara, CA, USA). To ensure robustness and protect against over-fitting 5-fold cross validation was used throughout. Data was split by classes into 5 equal parts and 5 ANNs were trained using an 80/20 percent training/validation split.

All models consisted of an input layer with 4374 nodes and a drop-out rate of 0.5 (to account for technical variability in single-cell data). Because we were interested in the shared logic between the species, rather than gene expression kinetics, expression levels were binarized (1 if the gene was expressed at any level, and zero otherwise) prior to learning. A 16-unit hidden layer with ReLU activation and L1 regularization (l=0.001), and a 14-unit softmax output layer. Training was performed for 21 epochs with a step size of 42, and a sample generator to re-sample 5 training examples per class per step. Loss was calculated using cross-entropy and gradient descent optimization was conducted using RMSprop with default parameters. For training in the target domain, the step size was set to be proportional to the number of training examples up to a maximum of 30 steps per epoch to avoid overfitting. Each model from the source domain was re-trained on 1, 2, …, 10, 15, 20, 25, 30 examples per class (excluding unrepresented classes in the target domain) using 5-fold cross validation.

### Evaluation of classification performance

To account for the extreme class imbalance, balanced accuracy (BA)^40^ was adapted from the binary setting described in Brodersen et al. 2010 to the multi-class setting. BA was calculated as the arithmetic mean of sensitivity (true positive rate) and specificity (true negative rate) for a given class against all other classes. Overall performance across all classes was calculated as the average balanced accuracy. Further, the F1 score was calculated from the harmonic mean of sensitivity and precision (predicted positive rate). Performance metrics were reported as the average from 5-fold cross validation. For analyses related to Fig. 2, classification performance of the source ANN in target domain was calculated from an ensemble of all 5 ANNs via plurality vote. Performance in the target domain was independently assessed using a test set containing all cell not used in training (~95% of human data; range: 17.2% to 98.7% of human cells per class).

### Topology of misclassification

The relationship between classes is specified by an acyclic graph **G**, the cell lineage tree (see Fig. 1d for reference), which is a hierarchical representation of the developmental history of cell types that is derived elsewhere (see for instance Tusi et al. 2018). In this graph, cell types are nodes and edges are direct developmental trajectories. We consider cells to be correctly classified, if the distance between the true label and the predicted label along the edges in **G** is *d* = 0. Misclassification events are categorized as proximal if the distance between true and predicted labels *d* = 1; and distal if the distance between labels *d* > 1.

### Sensitivity analysis

Because we binarized data prior to learning we were able to systematically assess the mutual information between input gene expression patterns and out posterior probabilities from the softmax output layer. Specifically, the mutual information^41^ was calculated using the input and output of the ANN trained on mouse data. To do so, the discretized gene expression ***x***_g_ ∈ {0,1} for gene g and the discretized posterior probabilities ***y***_c_ ∈ {[0, 1/3], (1/3, 2/3], (2/3, 1]} for class c were extracted from the training data and cross tabulated to obtain an ***x***_g_ by ***y***_c_ contingency matrix M. By normalizing M by the total number of observations, an estimate of the joint probability distribution 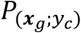, as well as estimates of the marginal distributions 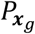 and 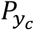 were obtained, from which the mutual information

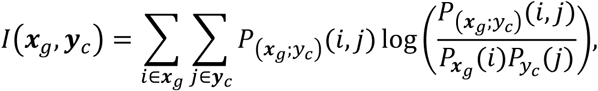

was calculated. Genes were then ranked for each class based upon mutual information, as a measure of the extent to which they contribute to class assignments.

### Gene set analysis

To obtain a functional annotation of the ranked gene lists obtained from the sensitivity analysis (see above), gene set analysis was performed, using the 100 highest ranking genes for each class as an input to the functional annotation tool in DAVID^42^ (v6.8; https://david.ncifcrf.gov/) and reference gene sets defined in the *biological process* gene ontology (GO).

### Data availability

Data reported in this work are available from ArrayExpress under accession E-MTAB-8629 and E-MTAB-8630.

## Acknowledgements

We would like to thank Alistair Bailey (University of Southampton) for his helpful discussion of Keras. This research was funded by the Medical Research Council (MC_PC_15078) and the Research Management Committee at the University of Southampton, Faculty of Medicine. ROCO acknowledges support from the UK Regenerative Medicine Platform “Acellular / Smart Materials – 3D Architecture” (MR/R015651/1), the Rosetrees Trust, Wessex Medical Research and the Biotechnology and Biological Sciences Research Council (BB/P017711/1).

## Author Contributions

Conceptualization: PSS, FA and BDM; Methodology: PSS, JJW, MRZ, RCGS, FA, YK and BDM; Investigation: PSS, TN, HI, YK, YS and FA; Data Curation: PSS; Formal Analyses: PSS, DD; Visualization, PSS and BDM; Supervision: BDM and MN; Writing – Original Draft: PSS, BDM; Writing – Review and Editing: all authors; Funding Acquisition: BDM, FA, KA, JJW, PSS; Infrastructure, Resources: FA, KA, ROCO, BDM; Project Administration: PSS, BDM.

## Disclosure of Conflicts of Interest

The authors declare that the research was conducted in the absence of any commercial or financial relationships that could be construed as a potential conflict of interest.

---

**Supplementary Figure 1.**
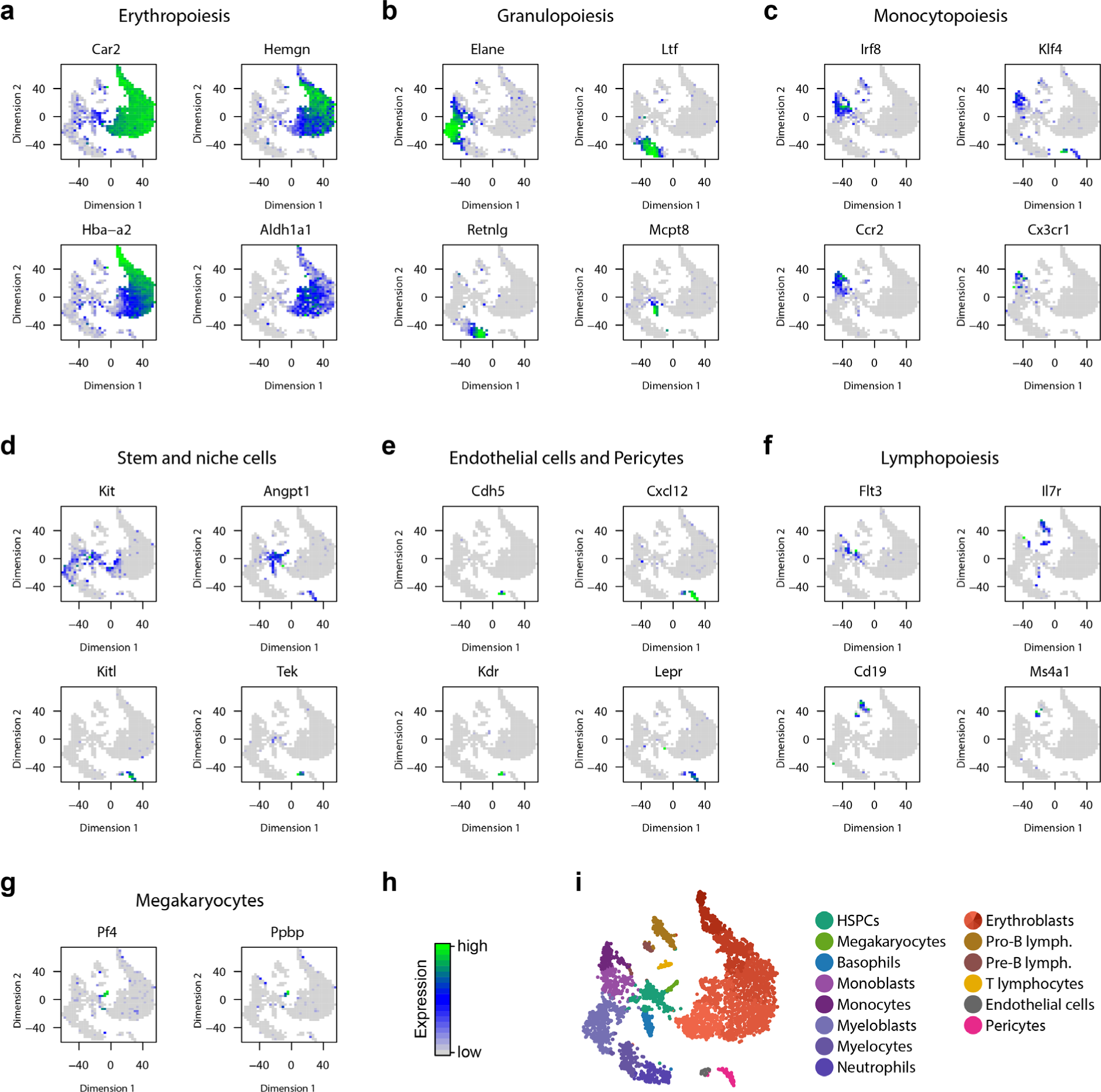
Gene expression localizes to developmental branches of mouse hematopoiesis. **a-g** Average gene expression superimposed onto 2D embedding of scRNAseq data. Displayed are the mean expression values for each 2D-bin. Localized expression indicative of **a** erythropoiesis, **b** granulopoiesis, **c** monocytopoiesis, **d** hematopoietic stem and progenitor cells and niche cells, **e** endothelial cells and pericytes, **f** lymphopoiesis, **g** thrombopoiesis. **h** Color scale. **i** Cluster structure from Fig. 1c for reference.

**Supplementary Figure 2.**
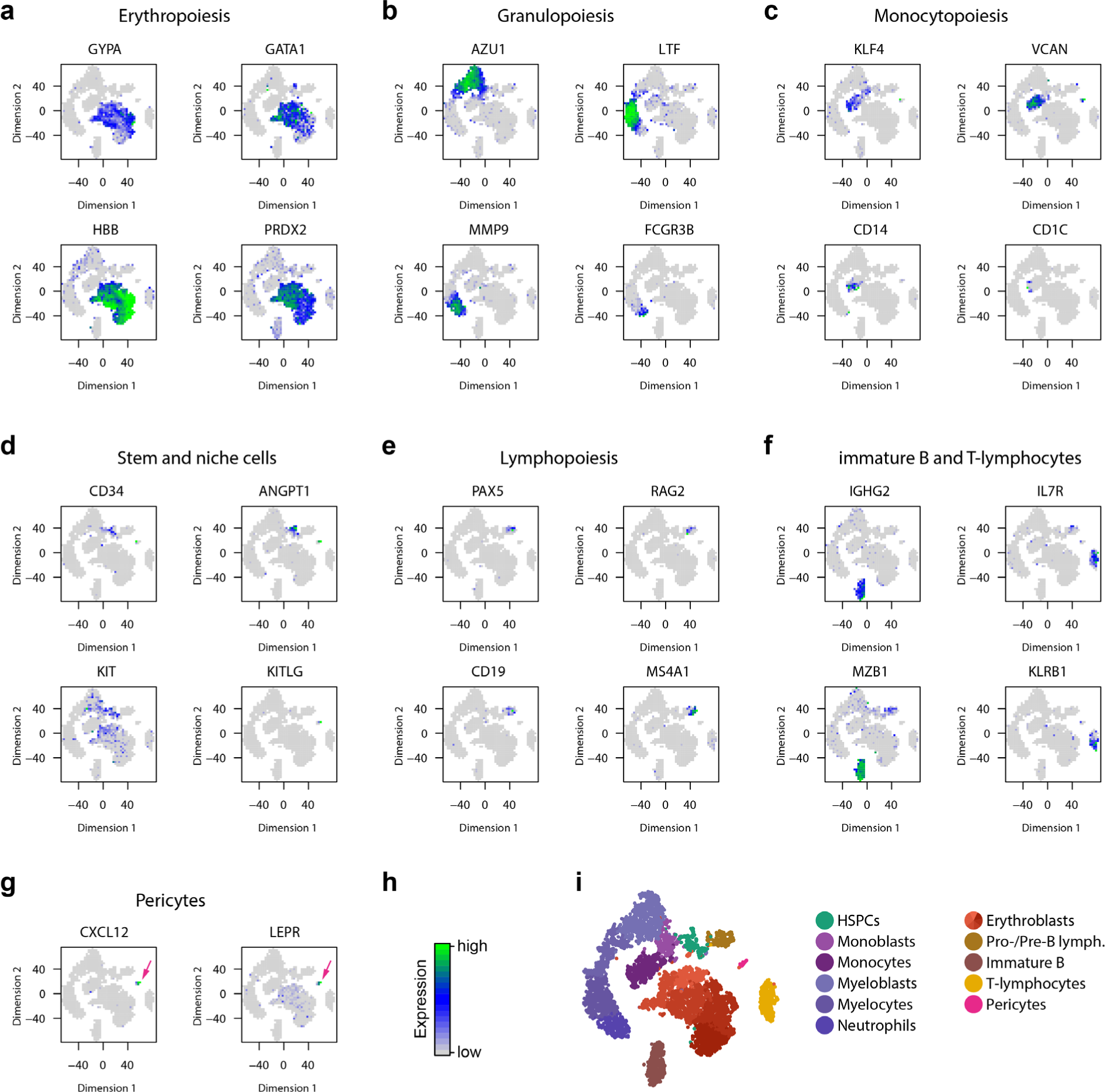
Gene expression localizes to developmental branches of human hematopoiesis. **a-g** Average gene expression superimposed onto 2D embedding of scRNAseq data. Localized expression indicative of **a** erythropoiesis, **b** granulopoiesis, **c** monocytopoiesis, **d** hematopoietic stem and progenitor cells and niche cells, **e** endothelial cells and pericytes, **f** lymphopoiesis, **g** pericytes. **h** Color scale. **i** Unsupervised clustering and annotation derived from the literature.

**Supplementary Figure 3.**
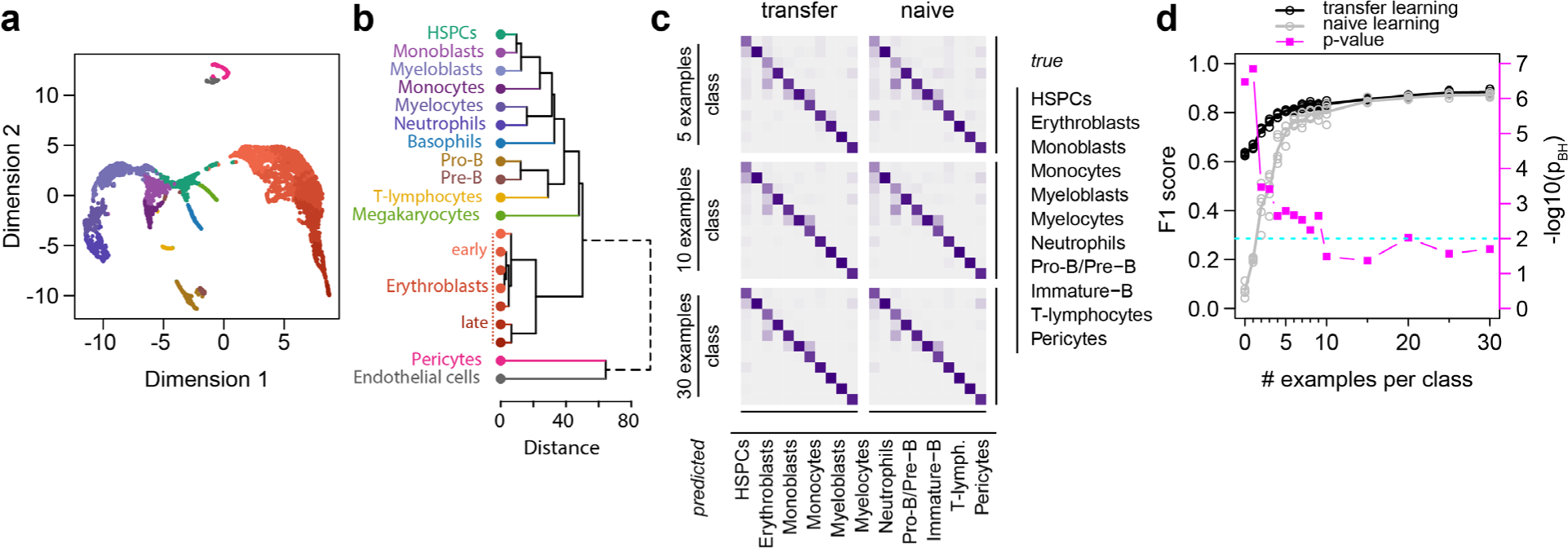
**(a)** Related to Fig. 1b, c. Alternative projection of mouse bone marrow scRNAseq data onto two dimensions using UMAP^43^. Data points represent cells, colored by cluster membership detailed in Fig. 1. **(b)** Related to Fig. 1b-e. Dendrogram of mouse bone marrow cell type dissimilarity. Displayed is the Euclidean distance between cluster median-centres^44^, calculated from the first 11 principal components. **(c)** Related to Fig 3. Confusion matrices at various levels of re-training (5, 10, 30 examples per class) for transfer learning and naïve learning. **(d)** Related to Fig 3. Average F1 score over all classes from 5-fold cross validation (primary *y*-axis), and corresponding negative logarithm (base 10) of *p*-values (FDR corrected; secondary *y*-axis), from one-tailed paired t-tests (alternative: F1_transfer_ > F1_naive_). Dashed line denotes a significance level of α = 0.01.

## Supplementary Material

### Human bone marrow cell characterization

To enrich the progenitor and niche cell subsets contained in bone marrow mononuclear cells (BM-MNCs), magnetic cell sorting (MACS) was employed to deplete cells expressing pan-leucocyte marker CD45 [*PTPRC*] or erythrocyte marker CD235a [*GYPA*] as well as to enrich skeletal stem cell marker STRO-1 [*HSPA8*].^45^ To discriminate broad classes of cells among the BM-MNCs, unsupervised clustering was employed at low resolution in Seurat (resolution parameter = 1.1; see Fig. S2i). This revealed the presence of 16 distinct cell types (including five erythroblast clusters and two myelocyte clusters that were each summarized as one cluster each, due to the apparent homogeneity analogous to mouse erythroblasts, compare Fig. S3b). At this deliberately low resolution, individual genes possessed sufficient discriminative power for their identification (see Fig. S2a-g). For instance, specific stages of neutrophil development can be identified based on the enzyme content of the secretory vesicles,^46^ such as primary azurophilic granules (*AZU1*), secondary specific granules (*LTF*), and tertiary gelatinase granules (*MMP9*), while mature neutrophils are identified based on the characteristic expression of CD16 (*FCGR3B*; Fig. S1b). Moreover, CD14 positive monocytes and CD1C positive dendritic cells^47^ can be identified among monoblasts expressing versican (*VCAN*; Fig. S2c). Additionally, the BM-MNC population contains a number of hematopoietic stem and progenitor cells (HSPCs), marked by the surface antigen CD34, *KIT*, and *ANGPT1* (Fig. S2d). Notably, a small but distinct subset of cells is marked by high levels of *CXCL12* (Fig. S1g), an important hematopoietic niche factor that, in mouse, is secreted by both osteoblasts at the endosteal surface^30^ and by pericytes at the endothelial interface^48^, and expression of Leptin receptor (*LEPR*), another marker of pericytes and adipocytes.^28^ As another example, lymphocytes such as Pro-B- and Pre-B-lymphocytes characterized by CD19 and CD20 (*MS4A1*) respectively can be distinguished from more mature B-lymphocytes marked by IgG heavy chain (*IGHG2*) and *MZB1* (a co-chaperone important for immunoglobulin-folding^49^) respectively (Fig. S2e-f).

### Modification to the bioinformatics pipeline to resolve multiple alignments

See separate document.

